# Nucleus accumbens sub-regions experience distinct dopamine release responses following acute and chronic morphine exposure

**DOI:** 10.1101/2024.06.28.601282

**Authors:** Sarah Warren Gooding, Elinor Lewis, Christine Chau, Suhail Sandhu, Julianna Glienke, Jennifer L. Whistler

## Abstract

It is well established that dopamine neurons of the ventral tegmental area (VTA) play a critical role in reward and aversion as well as pathologies including drug dependence and addiction. The distinct effects of acute and chronic opioid exposure have been previously characterized at VTA synapses. Recent work suggests that distinct VTA projections that target the medial and lateral shell of the nucleus accumbens (NAc), may play opposing roles in modulating behavior. It is possible that these two anatomically and functionally distinct pathways also have disparate roles in opioid reward, tolerance, and withdrawal in the brain. In this study we monitored dopamine release in the medial or lateral shell of the NAc of male mice during a week-long morphine treatment paradigm. We measured dopamine release in response to an intravenous morphine injection both acutely and following a week of repeated morphine. We also measured dopamine in response to a naloxone injection both prior to and following repeated morphine treatment. Morphine induced a transient increase in dopamine in the medial NAc shell that was much larger than the slower rise observed in the lateral shell. Surprisingly, chronic morphine treatment induced a sensitization of the medial dopamine response to morphine that opposed a diminished response observed in the saline-treated control group. This study expands on our current understanding of the medial NAc shell as hub of opioid-induced dopamine fluctuation. It also highlights the need for future opioid studies to appreciate the heterogeneity of dopamine neurons.

**Significance Statement:** The social and economic consequences of the opioid epidemic are tragic and far-reaching. Yet, opioids are indisputably necessary in clinical settings where they remain the most useful treatment for severe pain. To combat this crisis, we must improve our understanding of opioid function in the brain, particularly the neural mechanisms that underlie opioid dependence and addictive behaviors. This study uses fiber photometry to examine dopamine changes that occur in response to repeated morphine, and morphine withdrawal, at multiple stages of a longitudinal opioid-dependence paradigm. We reveal key differences in how dopamine levels respond to opioid administration in distinct sub-regions of the ventral striatum and lay a foundation for future opioid research that appreciates our contemporary understanding of the dopamine system.

## Introduction

In the last two decades, opioid overdose has devastated communities across America, causing massive healthcare burdens and claiming hundreds of thousands of lives. In 2021 there were 220 opioid overdose deaths per day, accounting for more than 75% of all drug overdose deaths in the United States (Spencer et al., 2022). Opioid overdose and associated risk behaviors tightly parallel the related phenomena of opioid dependence, addiction, and opioid use disorder (OUD). Opioid use behaviors have long been associated with changes in the activity of an opioid-sensitive mesolimbic dopamine circuit.

Activation of the mu-opioid receptor (MOR) by opioid agonists reduces GABA release on dopamine neurons in the VTA (Johnson and North, 1992; Jalabert et al., 2011; Matsui and Williams, 2011; Matsui et al., 2014). The mechanism of acute opioid reward is thought to rely on this disinhibition phenomenon leading to increased dopamine release in target regions, particularly the nucleus accumbens (NAc) (Ikemoto, 2007; Fields and Margolis, 2015). Repeated activation of this mesolimbic pathway by opioids leads to long-lasting plasticity in both VTA dopamine neurons and their downstream targets (Luscher, 2016). An overall decrease in dopamine function is traditionally understood as a hallmark of tolerance and withdrawal from opioids and other drugs of abuse (Koob et al., 1998). Chronic opioid exposure changes the opioid-sensitivity of presynaptic GABA neurons in the midbrain (Madhavan et al., 2010a; Matsui et al., 2014) and blockade of MOR signaling following chronic activation produces a strong increase in GABA release onto dopamine neurons in slice (Bonci and Williams, 1997; Madhavan et al., 2010b). However, few studies to date have examined the effects of repeated morphine treatment on dopamine release (Acquas et al., 1991; Pothos et al., 1991; Acquas and Di Chiara, 1992; Spanagel et al., 1993; Mazei-Robison et al., 2011; Lefevre et al., 2020). These studies vary in experimental design elements such as probe location, stimuli used, drug administration schedule, and control strategy. Such variation makes it challenging to synthesize these results into a cohesive model of how dopamine release is impacted by chronic opioid treatment.

The cellular composition of the VTA is rich in phenotypic diversity and a variety of cell-types have been genetically and functionally identified (Lammel et al., 2008; Pupe and Wallen-Mackenzie, 2015; Morales and Margolis, 2017). VTA dopamine neurons project to several distinct regions including the NAc (Beier et al., 2015), and NAc sub-regions receive dopaminergic input from anatomically distinct sections within the VTA differently distributed along medial-lateral and rostral-caudal axes depending on their NAc targets (Ikemoto, 2007). In addition to this topographical organization, VTA dopamine neurons show potentiation under different reward and aversion conditions depending on their projection target (Lammel et al., 2011) and also promote opposing behavioral responses (Lammel et al., 2012). Not all dopamine neurons in the VTA react the same way to rewarding and aversive stimuli (Lammel et al., 2014) and recent evidence suggests important differences in how projections to separate NAc sub-nuclei respond under rewarding versus aversive conditions (de Jong et al., 2019; Yuan et al., 2019). The prior physiological studies of opioid and opioid withdrawal mechanisms have not examined whether there is a distinction in opioid effect based on the projection targets of individual dopamine neurons. Some studies have implicated the medial portion of the NAc as the primary locus of drug reward (Fenu et al., 2006; Ikemoto, 2007; Corre et al., 2018), but little clarification has been made for the mechanisms that are altered following repeated drug use such as tolerance and dependence. Specifically, it is unclear if opioid reward and the aversive state of opioid withdrawal result from bidirectional modulation of the same circuit, or anatomically separable activities. Here we use a longitudinal fiber photometry paradigm to examine the acute and chronic impacts of morphine on dopamine release in the medial and lateral shell of the NAc.

## Methods

### Mice

Male C57BL/6 mice were purchased from The Jackson Laboratory (Sacramento, CA). Mice acclimated to the housing conditions for at least 5 days after arriving and were housed in groups of 2-5under a standard 12/12-h light/dark cycle, with ambient temperature set at 20 °C to 22 °C and food and water ad libitum. Initial surgeries were performed on mice aged 8–9 weeks and photometry experiments began at ∼13 weeks of age.

### Drug Preparation

All drugs were prepared in sterile physiological saline for injection volumes of 10 mL/kg (300 uL for a 30 gram mouse). Morphine sulfate (Mallinckrodt Pharmaceuticals) and naloxone hydrochloride (Sigma Aldrich) were prepared fresh from powder and used within seven days of dissolution. Brevital sodium (Par Pharmaceutical) was prepared fresh on day of use from 25 mg/mL stock aliquots stored at -20 °C. Veterinary carprofen (Rimadyl®) was diluted from 50 mg/mL sterile injectable solution to 0.5 mg/mL and used within 30 days of dilution.

### Stereotaxic Surgeries

Mice were anesthetized with inhaled 2% isoflurane in oxygen and immobilized in a stereotaxic frame (Model 1900, Kopf Instruments) and injected with 100nL of AAV1-hSyn-dLight1.3b (Patriarchi et al., 2018) in either the medial (bregma: 1.50, lateral: 0.65, ventral: -4.75) or lateral (bregma: 1.00, lateral: 2.00, ventral: -4.50) shell of the nucleus accumbens. Virus was prepared by the UC Davis Molecular Construct and Packaging Core with a titer of ∼10^14^ infectious units/mL. Virus was injected with a 33g stainless steel needle coupled to a 1 uL Hamilton syringe with polyethylene tubing backfilled with mineral oil at a rate of ∼100nL/min with a microsyringe pump (World Precision Instruments). The needle was withdrawn slowly five minutes following the completion of virus infusion. Following virus infusion, a borosilicate optic fiber (400 um, NA: 0.66, Doric Lenses), was implanted at the injection site and secured to the skull with Metabond dental cement (Parkell, Inc.) and a cement headcap was created to close the incision. At the time of implantation, a custom laser cut stainless steel headplate (manufactured by TEAM Prototyping Lab at UC Davis, design courtesy of the Stephan Lammel lab at UC Berkeley) was attached to the headcap to facilitate head-fixed photometry recordings. Mice were given carprofen (5 mg/kg, s.c.) at the time of surgery and every 24 hours for three days as a post-operative analgesic. No opioid analgesics were given until the time of photometry experiments to keep animals opioid-naïve.

### Jugular Vein Catheterization

Four weeks following viral injection and fiber implantation, mice were anesthetized with inhaled 2% isoflurane in oxygen. A round tipped polyurethane catheter (Instech Labs) was inserted into the left jugular vein, trimmed to appropriate length for body size and attached to a compatible vascular access button (Instech Labs) that was then implanted subcutaneously between the shoulder blades. Mice were given carprofen (5 mg/kg, s.c.) at the time of surgery and every 24 hours for three days as a post-operative analgesic.

Immediately following surgery, jugular catheters were flushed with physiological saline. Additional saline flushes were given every ∼3 days and following i.v. drug injections to clear residual drug from the catheter. Mice were allowed to recover for 4-7 days prior to first photometry recordings. 4-24 hours prior to first photometry recording, catheter patency was tested with a bolus of brevital sodium 25 mg/kg i.v. (less if mouse was unconscious before completion of injection). Mice that were mobile within 5 seconds of injection bolus were removed from the study. Following the final photometry recording, catheter patency was tested again. Mice were excluded from longitudinal analyses if catheter patency was lost before the conclusion of photometry experiments.

### Fiber Photometry

For all fiber photometry recordings, mice were placed on a freely-rotating running wheel and head-fixed. Fiber photometry recordings were collected with a commercial fiber photometry system (Doric Lenses) that transmits LED light at 465 nm sinusoidally modulated at ∼209 Hz and 405 nm sinusoidally modulated at ∼308 Hz through a fluorescence mini-cube and into a fiber-optic cable that is coupled to the implanted fiber cannula on the mouse. Fluorescent emission travels back through the emission filter in the mini-cube and is directed onto a photoreceiver and then converted to digital signal that is recorded by the Doric software. The two channels are demodulated to separate dopamine and control (isosbestic) signals and decimated to 120 Hz for recording to disk. For each recording, 15 minutes of baseline was collected prior to i.v. injection and the recording continued for 15 minutes post-injection. Recording sessions that included two injections were completed in the same manner with the mouse remaining on the recording apparatus and a new baseline period beginning 15 minutes after the first injection.

### Photometry Analysis

Preprocessing and analysis of photometry data was conducted using custom R scripts. The first five minutes of baseline recording were removed for plotting and peaks analysis. To calculate dF/F, a simple linear model was used to regress values from the 465 nm channel onto the 405 nm values during baseline period of the recording (before drug injection). This model was then used to predict 465 nm values based on the 405 nm values for the entire recording. The dF/F was defined as (465 value – predicted 465 value)/predicted 465 value. dF/F signals were converted to z-scores using the mean and standard deviation from the baseline period: z-score = (dF/F – baseline mean)/baseline SD. A thirty second rolling average was applied to the z-scores to smooth data for group analysis. For peak analysis, data was pre-processed using a method adapted from Martianova et al. (Martianova et al., 2019). Briefly, an adaptive iteratively reweighted penalized least squares (airPLS) algorithm is used to dynamically detect the slope of each channel so it can be subtracted for a flattened trace that maintains high frequency signal changes. These adjusted signals are then standardized using the median and standard deviation of the baseline period: standardized signal = (signal – baseline median)/baseline SD. A robust linear model is used to predict the standardized 465 signal based on the standardized 405 and dF/F is calculated as the actual standardized 465 nm values minus the predicted values. Peak detection was performed on the standardized dF/F by calculating a rolling threshold defined as 2.91 times the median absolute deviation (threshold chosen based on (Calipari et al., 2016)) of a 30 second rolling window. Local maxima are then detected using the findpeaks() function from the pracma package with a minimum distance of 0.5 seconds between peaks. Local maxima that fall below the rolling threshold are considered background signal and excluded.

### Daily Injections

Daily subcutaneous injections of morphine (10 mg/kg) or saline were given in the home cage between the hours of 10AM and 2PM. No subcutaneous injections were given on days when photometry recordings occurred.

### Histology

Following the conclusion of the study, mice were anesthetized with isoflurane and transcardial perfusion was performed with 4% paraformaldehyde. Brains were extracted and post-fixed in 4% PFA for minimum 24 hours. 60 um coronal sections were cut and incubated overnight in a primary antibody solution with polyclonal chicken anti-GFP (ab13970, Abcam). The following day, slices were washed and stained for 1-4 hours with goat anti-chicken Alexa Fluor 488 (A-11039, Invitrogen) secondary solution. Sections were mounted with glass coverslips using Vectashield Hardset with DAPI. Images were acquired using a Zeiss LSM 800 confocal microscope.

### Experimental Design and Statistical Analyses

Statistics were conducted in R with the RStudio software. Comparison of baseline spontaneous frequency and amplitude was done with unpaired two sample t tests. For comparison of spontaneous transient data following morphine or saline injection, a two-way ANOVA was used to examine simple main effects and interaction effects between drug injection (morphine vs saline) and recording location (medial vs lateral NAc shell). Darly and late components of the recording were analyzed separately. To compare the effects of chronic treatment, two-way ANOVA with repeated measures was used with chronic treatment condition (morphine vs saline) as a between-subject factor and recording day (Day 0 vs Day 7) as a within-subject factor. If significant main effects were found, post-hoc analyses were conducted using Tukey’s multiple comparisons test.

### Code Accessibility

Custom scripts used for data processing and peaks analysis are available at Github.

## Results

### Morphine injection induces anatomically distinct dopamine increases in the medial and lateral NAc

To compare the effects of morphine on dopamine release in the medial and lateral shell of the nucleus accumbens, we used an *in vivo* fiber photometry paradigm with IV morphine injection (Fig. 1A). Mice were injected with virus carrying the genetically encoded dopamine sensor dLight 1.3b in either the medial or lateral shell of the nucleus accumbens and a fiber optic cannula was implanted at the injection site. Four weeks after viral injection and fiber implantation, mice were implanted with jugular vein catheters to allow for instantaneous delivery of drug. Head-fixed fiber photometry recordings consisted of a 10-minute baseline period followed by an IV bolus of morphine (10 mg/kg) and an additional 15 minutes of recording. Individual subject recordings reveal both fast (sub-second) and slow (several minute) dopamine transient activity (Fig. 1B). Morphine injection produced a large transient increase in overall dopamine signal in the medial shell of the nucleus accumbens characterized by a steep increase of the moving average followed by a short decay with a plateau above baseline after several minutes (Fig. 1C, purple). In the lateral shell, minimal increase was observed in response to IV morphine with any increase appearing to ramp up over the 15-minute recording period (Fig. 1C, blue). Heatmaps of individual mice are shown in Fig. 1D.

**Figure 1:**
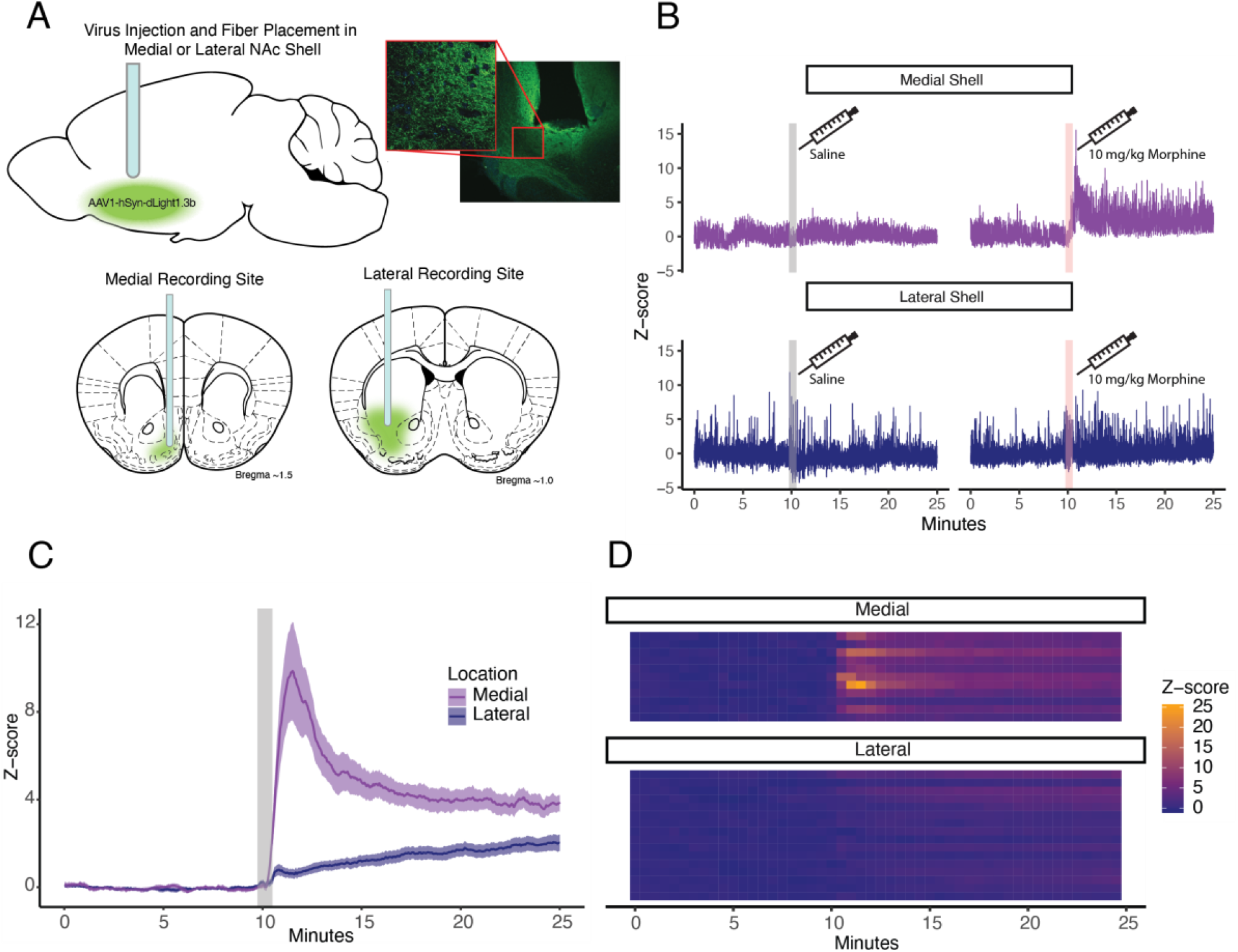
Morphine injection induces anatomically distinct dopamine increases in the medial and lateral NAc. A) Schematic of fiber photometry strategy. Optic fibers were implanted at the time of viral injection to allow expression of dLight1.3b in either the medial or lateral shell of the NAc. Approximate injection sites and fiber locations are shown in coronal sections. B) Representative traces of dopamine signal at baseline and following IV injection of morphine (right) or saline (left). C) Average dopamine response to IV morphine injection in the medial and lateral shell of the nucleus accumbens. Data has been smoothed over a 30 second rolling window. Shaded region reflects SEM. D) Heatmaps showing individual subject responses to IV morphine in 30 second time bins. Each row is a recording session for one mouse.

### Morphine injection alters spontaneous dopamine transient activity

To examine the faster sub-second dopamine transients (Fig. 2A), we used a different pre-processing algorithm to dynamically remove the slope of the recording (Martianova et al., 2019) and then performed peak analysis to identify high pass dopamine transient events reflecting spontaneous dopamine activity. We first examined the patterns of peak count and peak prominence, defined as true peak height minus the local moving average, across the entire recording by visualizing them in one minute time bins (Fig. 2B) to see if the time course of peak changes followed that of the larger but slower, morphine-induced dopamine changes (Fig. 1C). We found that fluctuations in peak prominence and frequency had a similar time course to the large smoothed dopamine signal. Based on this time course, we divided the recordings into two distinct epochs for further analysis. The data was binned into the early response component, defined as the five minutes directly following injection, and the late response component, which was the remaining ten minutes of the recording.

**Figure 2:**
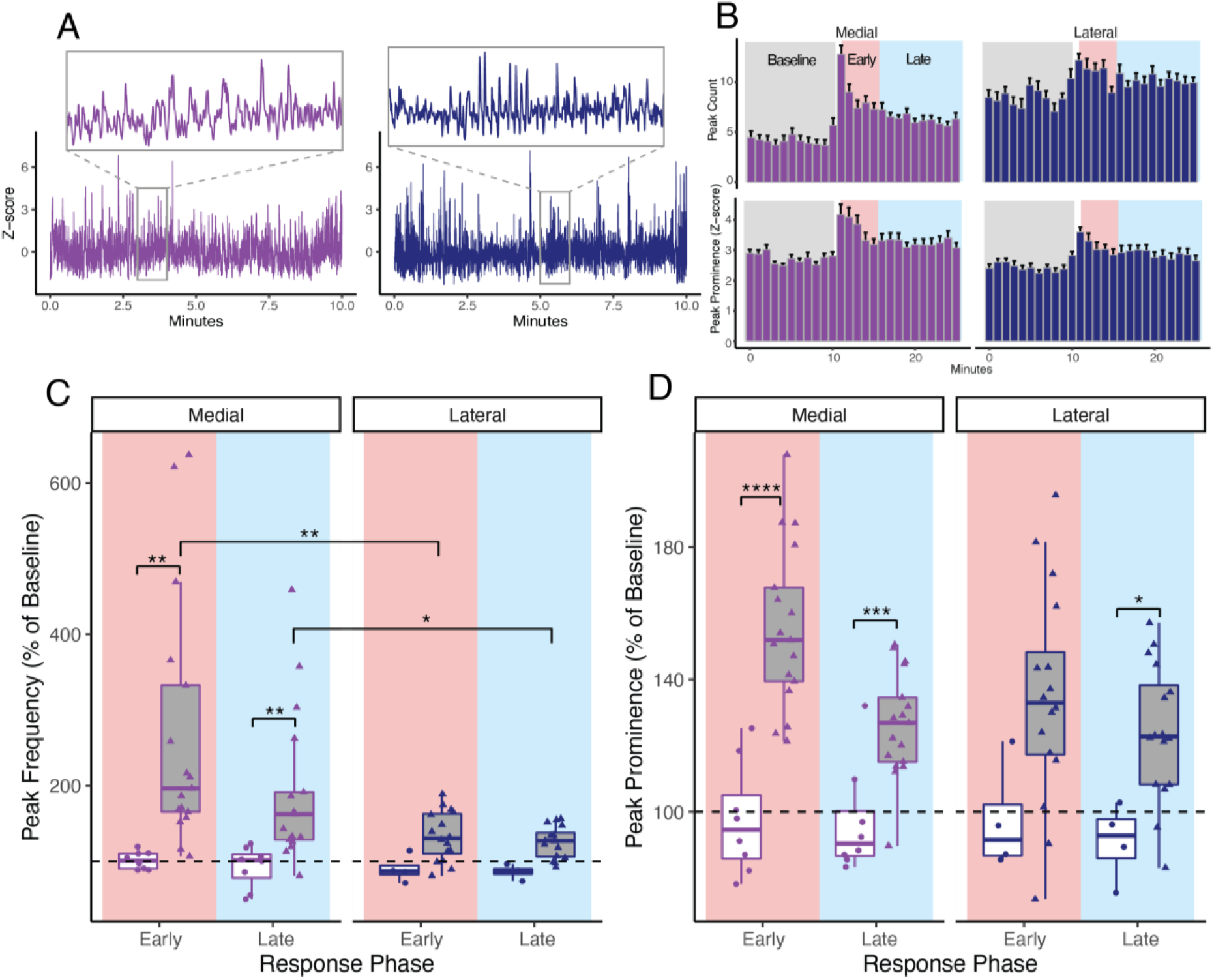
Morphine injection alters spontaneous dopamine transient activity. A) Representative traces showing spontaneous dopamine transient activity in the medial and lateral NAc shell at baseline. B) Peak count (top) and peak prominence (bottom) across the entire recording session. Data have been separated into one-minute bins to display average and standard error of each minute of the recording.C) Peak frequency in the medial and lateral NAc shell during early (red) component and late (blue) component response periods after IV injection of morphine or saline. Data is normalized to peak frequency during the pre-injection baseline period. Significant differences were found between medial morphine and saline response in early (p = 0.004) and late (p = 0.007) recording components and between the medial and lateral morphine response in the early (p = 0.005) and late (p = 0.04) component (Tukey’s multiple comparisons test after two-way ANOVA). D) Peak prominence in the medial and lateral NAc shell during early component and late component response periods after IV injection of morphine or saline. Data is normalized to peak prominence during the pre-injection baseline period. Significant differences were found between medial morphine and saline response in early and late recording components (p < 0.001 for both) and between lateral morphine and saline response in the late (p = 0.011) component (Tukey’s multiple comparisons test after two-way ANOVA). *p < 0.05, **p < 0.01, ***p < 0.001, ****p < 0.0001.

At baseline, there was a higher frequency of events in the lateral shell (0.149 peaks/s) than that of the medial shell (0.077 peaks/s) (t = 7.773, p < 0.001, Welch two sample t-test). The prominence of events was also slightly higher at baseline in the medial (2.707) than the lateral (2.435) NAc shell (t = -3.443, p = 0.001). Frequency of spontaneous events was increased by IV morphine injection in the medial but not the lateral shell (Fig. 2C). A two-way ANOVA found no significant interaction between the effects of implant location and morphine exposure during the early component (F_(1,44)_ = 2.402, p = 0.128) or the late component (F_(1,44)_ = 0.013, p = 0.909) response periods, but simple main effects analysis revealed a significant effect of morphine injection compared with saline in the early and late response components (p = 0.002 for both components). There was also a significant effect of implant location during early (p = 0.006) and late (p = 0.037) components. Event frequency in the medial shell was elevated (256% of baseline) directly following morphine injection, an effect that did not occur with IV saline injection (p = 0.004, Tukey’s multiple comparisons test). During the late component of the morphine response, spontaneous frequency remained elevated (184% of baseline) and significantly higher than the saline response (p = 0.007, Tukey’s multiple comparisons test) in the medial shell. This frequency increase was not detected in the lateral shell during the early or late components of the response and Tukey’s multiple comparisons test revealed a significant difference in the effect of morphine injection on normalized frequency between the medial and lateral shell in both early (p = 0.005) and late (p = 0.04) response components.

Peak prominence was analyzed similarly to frequency (Fig. 2D). There was no interaction between the effect of implant location or treatment on the prominence of peaks in the early (F_(1,44)_ = 1.655, p = 0.205) or late (F_(1,44)_ = 0.013, p = 0.909) response components in a two-way ANOVA. Simple main effects showed an effect of morphine exposure in both response epochs (p < 0.001 for early and late components) but no significant effect of implant location. Peak prominence in the medial shell was significantly elevated by morphine injection (158% and 129% of baseline respectively in early and late components) compared to saline in both response epochs (p < 0.001, Tukey’s multiple comparisons test). In contrast to what was observed with spontaneous frequency, peak prominence in the lateral shell was increased relative to baseline by acute morphine injection, a change that reached significance relative to saline injection in the late response epoch (124% of baseline) (p = 0.011) but remained slightly below significance in the early component (135% of baseline) (p = 0.065).

### Recurrent morphine treatment does not shift dopamine response to IV morphine in the NAc shell

Chronic opioid treatment and opioid tolerance are hypothesized to reduce dopamine neuron activity based on changes in inhibitory pre-synaptic activity (Bonci and Williams, 1997; Madhavan et al., 2010a; Matsui et al., 2014). To test the effect of chronic morphine on NAc dopamine release, we integrated our photometry recording protocol into a daily morphine treatment paradigm in which mice received six days of daily morphine (10 mg/kg, SC) or saline injection flanked by photometry recordings of IV morphine responses before and after this daily treatment paradigm (Fig. 3A). There was no difference detected in the response to morphine on day 7 compared to that observed on day 0 in mice that received daily morphine. This was true in both the medial and lateral NAc shell (Fig. 3B,C, left). Unexpectedly, in the medial shell of mice that received daily saline injections, there was a decreased response to morphine injection on day 7 compared to day 0 (Fig. 3B, right). The implication of this result is that daily morphine treatment precludes the diminished response to the secondary morphine injection observed in the saline treated mice.

**Figure 3:**
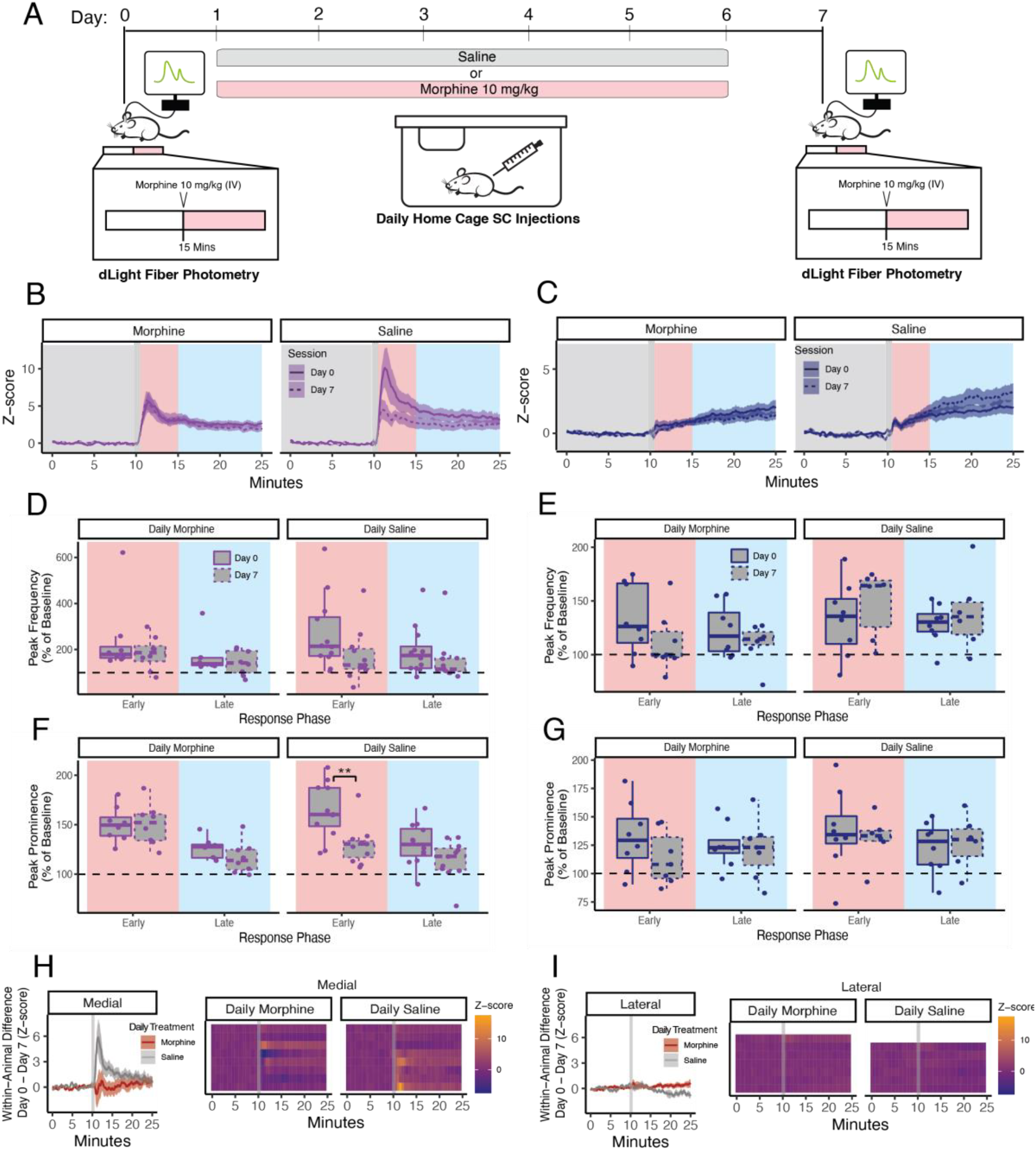
Recurrent morphine treatment does not shift dopamine response to IV morphine in the NAc shell. A) Timeline of longitudinal treatment paradigm. Mice received either morphine (10 mg/kg SC) or saline injection on days 1-6. On Days 0 and 7, identical fiber photometry recordings were conducted in which IV morphine (10 mg/kg) was injected 15 minutes into the baseline period. B) Average dopamine response to IV morphine injection in the medial shell of the nucleus accumbens on day 0 (solid line) and day 7 (dashed line). C) Average dopamine response to IV morphine injection in the lateral shell of the nucleus accumbens on day 0 (solid line) and day 7 (dashed line). D,E) Peak frequency in the medial (D) and lateral (E) NAc shell during early component and late component response periods after IV injection of morphine. Data is normalized to peak frequency during the pre-injection baseline period. F,G) Peak prominence in the medial (F) and lateral (G) NAc shell at baseline, early component and late component response periods before and after IV injection of morphine. Data is normalized to peak frequency during the pre-injection baseline period. There was a significant difference in the morphine response between day 0 and day 7 in mice treated with daily saline (p = 0.005, Tukey’s multiple comparisons test after two-way ANOVA with repeated measures). H,I) Within-subject differences in smoothed dopamine signal between day 0 and day 7 in mice treated daily with morphine (red) or saline (gray. Shown as group averages (left) and individual subject differences (right). Each heatmap row represents individual subject differences for one animal in 30 second time bins. **p < 0.01.

Peak analysis revealed comparable frequency of spontaneous events between day 7 and day 0 for both early and late response components in medial and lateral shell recordings in mice treated daily with morphine (Fig. 3D,E). A two-way ANOVA with repeated measures showed no significant interaction between recording day and daily treatment but there was a significant main effect of daily morphine treatment (p = 0.41). However, Tukey’s multiple comparisons testing revealed no significant differences between individual groups or between recording days. When we analyzed the effects of chronic drug treatment on peak prominence, we found a significant effect of drug treatment (p = 0.037, Two-way ANOVA with repeated measures) (Fig. 3F,G). Tukey’s multiple comparisons test revealed a significant decrease in peak prominence after morphine injection in the medial shell of mice who received daily saline treatment (p = 0.005). This decrease was limited to the early component of the morphine response, consistent with the diminished response seen in the smoothed data (Fig. 3B) where the height of the initial transient is reduced but the lingering morphine effect (>5 minutes post-injection) remains similar to that seen on day 0.

To confirm that group averaging was not masking a within-subject effect, we performed within-subject analysis of the morphine-induced bulk fluorescence effect (Fig. 3H,I). For each mouse, we subtracted the standardized fluorescence from day 7 from that of day 0 across the recording period. Like the grouped analysis this revealed a diminished dopamine response to IV morphine on the seventh day in the medial shell of mice that had received daily saline treatment peaking in the first 5 minutes post-injection. No other groups showed a strong effect of repeated morphine or saline treatment.

### Morphine withdrawal is not reflected by a dopamine release change in the medial or lateral NAc shell

Withdrawal from opioids in chronically treated animals is associated with increased inhibitory currents in dopamine neurons of the VTA (Bonci and Williams, 1997; Madhavan et al., 2010b) and there is some evidence of reduced dopamine release activity in the NAc following naloxone administration in opioid treated animals (Pothos et al., 1991). To compare the effect of naloxone-precipitated withdrawal on dopamine release in the medial and lateral shell of the NAc, we expanded the longitudinal model described above to capture a naloxone-induced dopamine response before and after daily morphine treatment (Fig. 4A). Briefly, mice received two initial fiber photometry recordings to capture their response to naloxone injection (5 mg/kg, IV) before and after six days of saline treatment. These initial recordings allowed us to observe the effect of naloxone on dopamine release in morphine naïve mice as well as to determine if this response changed across the timing of our morphine treatment paradigm with daily handling and saline injection alone. We found no deviation from baseline dopamine levels in response to naloxone injection beyond the small deflection that results from the injection artifact (Fig. 4B). This was the case for recordings on day -9 and day -2.

**Figure 4:**
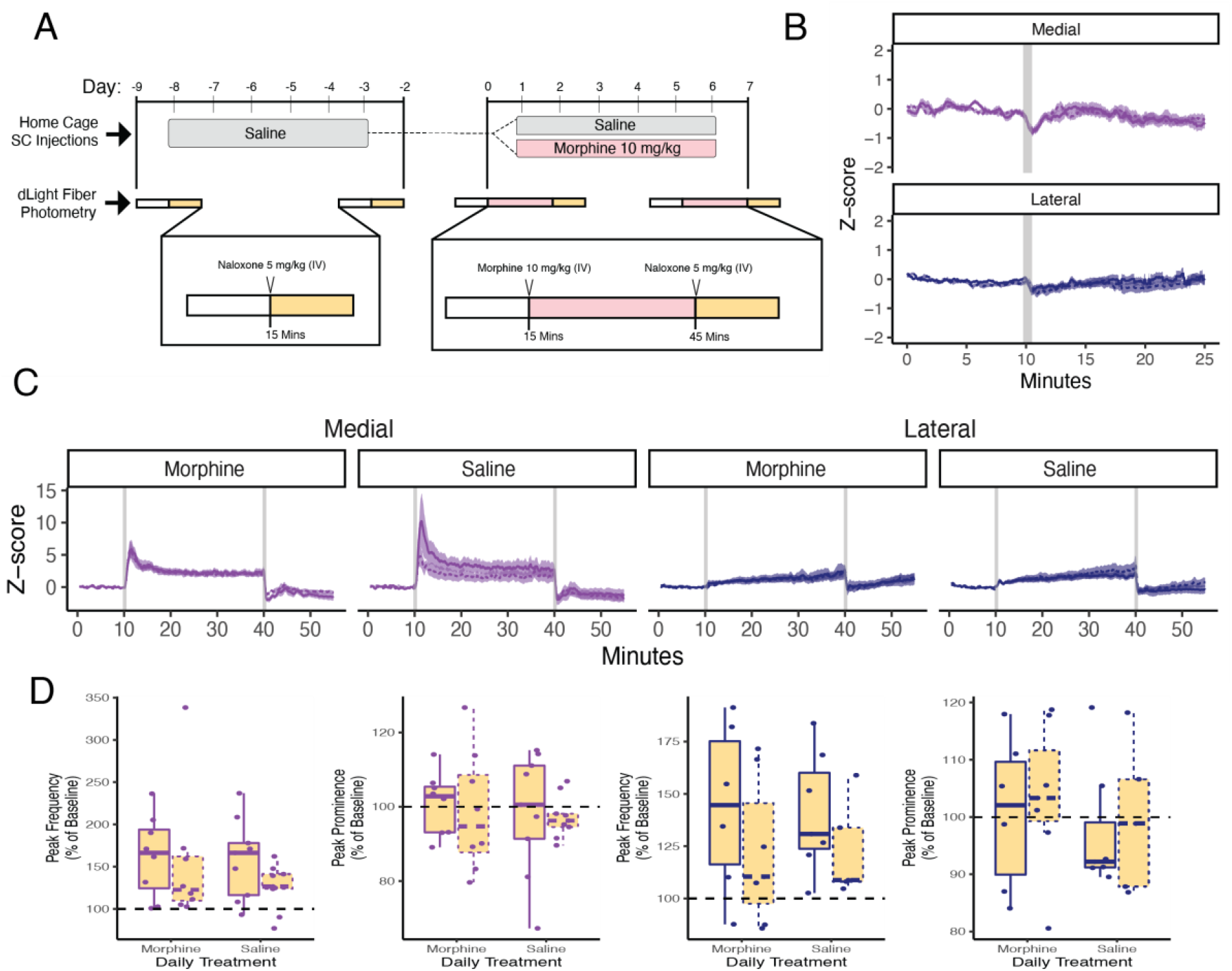
Morphine withdrawal is not reflected by a dopamine release change in the medial or lateral NAc shell. A) Timeline of morphine withdrawal paradigm. Morphine naïve mice received two fiber photometry (day -9 and day -2) recordings during which IV naloxone (5 mg/kg) was injected 15 minutes into the baseline period. These recordings were separated by six days on which mice received daily SC saline injections. Two days after the second naloxone recording (day 0), mice received fiber photometry recordings during which morphine was injected at 15 minutes and naloxone was injected at 45 minutes. Mice received either daily morphine or saline injection for the following six days and then a final fiber photometry recording on day 7 that matched the recording on day 0. B) Average dopamine response to IV naloxone injection in the medial and lateral shell of the NAc on day -9 (solid line) and day -2 (dashed line). C) Average dopamine response to IV morphine and subsequent naloxone injection in the medial and lateral shell of the NAc on day 0 (solid line) and day 7 (dashed line) in mice treated daily with morphine or saline. D) Peak prominence (top) and inter-peak interval (bottom) in the medial and lateral NAc shell after IV naloxone administration on day 0 (white) and day 7 (gray). E) Within-subject differences in smoothed dopamine signal between day 0 and day 7 in mice treated daily with morphine (red) or saline (gray).

Following these morphine naïve recordings, we performed the previously described recording paradigm to capture the response to IV morphine on day 0 and day 7 of the daily treatment schedule. On these recording days, mice were left on the recording apparatus where they received IV naloxone 30 minutes after their IV morphine injection. The goal of these recordings was to capture the dopaminergic changes that occur when morphine signaling is rapidly halted by naloxone in mice after acute morphine exposure (day 0) or following repeated morphine exposure (day 7). These IV naloxone injections in the presence of morphine produced a stark decrease in dopamine signal. This decrease was present in both the medial and the lateral shell, confirming that the slow rise in dopamine we observed in the lateral shell in response to morphine injection is an opioid-driven signal increase, despite being much more subtle than the increase observed in medial shell (Fig. 1C). While naloxone starkly reverses the elevated dopamine signal produced by IV morphine injection, we did not find any differences in the degree of naloxone-precipitated decrease between day 0 and day 7 in mice that were treated daily with morphine versus saline (Fig. 4C). Peak analysis of spontaneous dopamine transients following naloxone injection also showed no significant differences in peak frequency or prominence between day 0 and day 7 in morphine versus saline treated mice (Fig. 4D).

## Discussion

### Dopamine release is distinctly altered by morphine injection in the medial and lateral shell of the NAc

Here we examined dopamine release dynamics in the medial and lateral sub compartments of the NAc shell in response to intravenous morphine and morphine withdrawal. Our data align with the previous understanding that the medial shell is a hotspot for opioid-induced dopamine release. In the medial shell we observed significant dopamine changes in the minutes following morphine injection exemplified by bulk dopamine rise and increased frequency and amplitude of spontaneous events. We saw a more subdued response to IV morphine in the lateral shell of the NAc where the response had a much smaller magnitude with a slower time course. Interestingly, the amplitude of individual release events was significantly increased in the lateral shell minutes after morphine injection despite no change in frequency of these events.

While there is a history implicating the medial shell as a critical region for acute opioid reward and associated dopamine release (Cadoni and Di Chiara, 1999; Di Chiara, 2002; Gerrits et al., 2002; Fenu et al., 2006; Lecca et al., 2007; Bassareo et al., 2011; Bassareo et al., 2013), this oeuvre lacks similar characterization of the NAc lateral shell. This is salient given the dense dopaminergic projection from the VTA to this region that evidently does contain opioid-sensitive neurons, albeit fewer than found in the medial VTA (Corre et al., 2018). Given the extensive work comparing these two projections in the context of naturalistic rewarding and aversive stimuli (Lammel et al., 2011; Lammel et al., 2012; Lammel et al., 2014; de Jong et al., 2019; Yuan et al., 2019), we chose to explore the lateral projection in parallel with the medial projecting neurons in our opioid exposure model. While the response to morphine was far more dramatic in the medial shell, the lateral shell was not unaffected by acute morphine exposure. The slow-rising, naloxone-reversable dopamine response observed in the lateral shell, as well as the increased amplitude of spontaneous events make clear that this population should not be ignored.

### The mechanisms of opioid tolerance in the NAc still require extensive investigation

A key intention of this study was to characterize the effects of repeated opioid treatment on dopamine release in the medial and lateral NAc. Reduction of dopamine activity is commonly charged as a critical mechanism of morphine tolerance, but there are surprisingly few examples in the literature of the impact of chronic opioids on dopamine release. In this paradigm, six days of daily morphine treatment did not alter the dopaminergic effect of a morphine challenge injection in either of our recording sites. This occurred despite a dosing schedule we have found to be sufficient to induce both analgesic tolerance and morphine dependence (He et al., 2021). The existing literature on dopamine release in chronic opioid models is rife with contradictions and methodological caveats. Some work has found diminished responses that align with the traditional views of opioid tolerance (Mazei-Robison et al., 2011; McClain et al., 2023) while others have observed unchanged dopamine tone (Pothos et al., 1991) or even sensitization when responses in the NAc core are measured (Lefevre et al., 2020). The contrast of all these data highlights the ever-present trade-offs between mechanistic specificity and human relevance in our models and emphasizes the need to approach these questions with multiple experimental strategies.

The reduced response to morphine that we observed in the medial shell of mice following six days of saline treatment was entirely unexpected given that this effect was not present in the morphine-treated mice. The presence of an effect in the control group that was not observed with the experimental manipulation is perplexing, especially as care was taken to ensure that the groups were randomly assigned, housed together, and recordings were interspersed on the same days. It is important to note that these saline-treated mice were not morphine naïve on their final recording day, but rather, had received a single previous dose of morphine during their initial recording. It is therefore possible that a single IV morphine delivery is a strong enough stimulus and/or the photometry chamber and associated handling provides a distinct enough context to promote cue association that may shift future responses. Dopamine activity in the ventromedial shell does show a diminishing response to expected reward delivery compared to the strong response observed upon initial delivery (de Jong et al., 2019), an example of the reward prediction error concept. If the change in response we found in the saline control mice is driven by such a mechanism, our data would imply that chronic morphine treatment blocks this effect such that a morphine stimulus continues to elicit a strong transient dopamine response regardless of its expectation. This phenomenon, which we call “covert sensitization”, is not entirely new. There is work suggesting that drugs such as cocaine differ from natural reward sources by bypassing the classic reward prediction error effect (Keiflin and Janak, 2015). However, the paradigm used here was not intended to directly test the role of drug expectation, and to properly pursue this hypothesis would require adjustments to be made to the current design. Furthermore, our observation of this covert sensitization effect spotlights the importance of including both within-subject and between-subject controls in studies of chronic drug exposure.

### Unique challenges accompany the search for neural correlates of opioid withdrawal

While a dopamine decrease has been described previously during naloxone-precipitated withdrawal (Pothos et al., 1991), we employed a high temporal resolution method to capture the opioid withdrawal state *in vivo*. Our approach was also distinct in our choice to measure withdrawal shortly after morphine administration rather than following an abstinence period. There are reasonable arguments for and against this strategy as well as the alternative method of measuring a naloxone response relative to an abstinent baseline. Whether baseline dopamine concentration is effectively a moving target is a topic of central concern to this work. It is unclear how effectively basal dopamine levels can be compared across days using fiber photometry, and the literature is not settled regarding whether basal dopamine does (Acquas et al., 1991; Acquas and Di Chiara, 1992; Gerrits et al., 2002) or does not (Pothos et al., 1991; Spanagel et al., 1993) change over repeated morphine treatment. Given these caveats, we chose instead to examine how the shift from a morphine-adjusted baseline differed in a within-subject model where a naloxone response was measured before and after repeated morphine treatment.

While naloxone effectively reversed the increased dopamine tone that follows a morphine injection, this decrease was not influenced by six days of morphine treatment. This result was surprising as we hypothesized that, due to the known role of the nucleus accumbens in aversion, there would be a measurable effect in one or both of our recording sites that corresponded with the aversive state of naloxone-precipitated opioid withdrawal. Aversive behaviors and accompanying dopamine modulation depend on complex circuits upstream of dopamine neurons (Lammel et al., 2011; Lammel et al., 2012; Lammel et al., 2014; de Jong et al., 2019), so it is possible that the impact of disinhibiting opioid-sensitive presynaptic GABA neurons with an antagonist occludes modulation from these other afferent inputs. In this case, the aversion-related dopamine activity that accompanies drug withdrawal would be masked by a less subtle pharmacological effect.

Beyond the possibility that opioid withdrawal is not encoded in the regions we observed, there are several other factors that could have prevented us measuring a dependence-related morphine withdrawal response in dopamine release. Dopamine neurons are a mixed population which can be categorized in many ways beyond projection target (Garritsen et al., 2023). The spatial resolution of this extracellular recording method could be too low to resolve changes that occur within a subset of this richly heterogeneous population. Interpreting extracellular dopamine levels is inherently challenging given its influence by many factors beyond neural firing (de Jong et al., 2022). Volume transmission of dopamine leads to multiple timescales of transmission which blur the alignment of extracellular dopamine with cellular activity (Liu et al., 2021). Local circuitry in the nucleus accumbens such as cholinergic interneurons also regulate release from dopamine terminals (Cachope et al., 2012; Threlfell et al., 2012). Finally, expression levels of the dopamine transporter, can differ based on projection target (Lammel et al., 2008) and are likely altered by chronic drug treatment (Simantov, 1993) as well. Our results underscore the need for further investigation into the mechanisms underlying opioid tolerance and withdrawal, considering the intricate interplay of neural circuits and neurotransmitter systems within the NAc. As we continue to refine our understanding of dopamine modulation in opioid use, our findings emphasize the importance of employing multifaceted experimental approaches to unravel the complexities of opioid-induced neuroadaptations.

## Acknowledgements

This study was funded by the National Institutes of Drug Abuse under award numbers R01DA056543 (JLW), T32MH112507 (SWG), F31DA051116 (SWG), and by funds provided by the state of California as start up to JLW. The authors wish to thank the Stephan Lammel Lab at UC Berkeley for instruction and sharing of custom CAD files for components of the head fixing setup. We also wish to thank the UC Davis TEAM prototyping lab and Dr. Yi-Je Chen with the UC Davis microsurgery core for his training and technical assistance in the jugular catheter procedure.

## References

Acquas E, Di Chiara G (1992) Depression of mesolimbic dopamine transmission and sensitization to morphine during opiate abstinence. J Neurochem 58:1620–1625.

Acquas E, Carboni E, Di Chiara G (1991) Profound depression of mesolimbic dopamine release after morphine withdrawal in dependent rats. Eur J Pharmacol 193:133–134.

Bassareo V, Musio P, Di Chiara G (2011) Reciprocal responsiveness of nucleus accumbens shell and core dopamine to food- and drug-conditioned stimuli. Psychopharmacology (Berl) 214:687–697.

Bassareo V, Cucca F, Cadoni C, Musio P, Di Chiara G (2013) Differential influence of morphine sensitization on accumbens shell and core dopamine responses to morphine- and food-conditioned stimuli. Psychopharmacology (Berl) 225:697–706.

Beier KT, Steinberg EE, DeLoach KE, Xie S, Miyamichi K, Schwarz L, Gao XJ, Kremer EJ, Malenka RC, Luo L (2015) Circuit Architecture of VTA Dopamine Neurons Revealed by Systematic Input-Output Mapping. Cell 162:622–634.

Bonci A, Williams JT (1997) Increased Probability of GABA Release during Withdrawal from morphine. J Neurosci 17:796–803.

Cachope R, Mateo Y, Mathur BN, Irving J, Wang HL, Morales M, Lovinger DM, Cheer JF (2012) Selective activation of cholinergic interneurons enhances accumbal phasic dopamine release: setting the tone for reward processing. Cell Rep 2:33–41.

Cadoni C, Di Chiara G (1999) Reciprocal changes in dopamine responsiveness in the nucleus accumbens shell and core and in the dorsal caudate-putamen in rats sensitized to morphine. Neuroscience 90:447–455.

Calipari ES, Bagot RC, Purushothaman I, Davidson TJ, Yorgason JT, Pena CJ, Walker DM, Pirpinias ST, Guise KG, Ramakrishnan C, Deisseroth K, Nestler EJ (2016) In vivo imaging identifies temporal signature of D1 and D2 medium spiny neurons in cocaine reward. Proc Natl Acad Sci U S A 113:2726–2731.

Corre J, van Zessen R, Loureiro M, Patriarchi T, Tian L, Pascoli V, Luscher C (2018) Dopamine neurons projecting to medial shell of the nucleus accumbens drive heroin reinforcement. Elife 7.

de Jong JW, Fraser KM, Lammel S (2022) Mesoaccumbal Dopamine Heterogeneity: What Do Dopamine Firing and Release Have to Do with It? Annu Rev Neurosci 45:109–129.

de Jong JW, Afjei SA, Pollak Dorocic I, Peck JR, Liu C, Kim CK, Tian L, Deisseroth K, Lammel S (2019) A Neural Circuit Mechanism for Encoding Aversive Stimuli in the Mesolimbic Dopamine System. Neuron 101:133–151 e137.

Di Chiara G (2002) Nucleus accumbens shell and core dopamine: differential role in behavior and addiction. Behav Brain Res 137:75–114.

Fenu S, Spina L, Rivas E, Longoni R, Di Chiara G (2006) Morphine-conditioned single-trial place preference: role of nucleus accumbens shell dopamine receptors in acquisition, but not expression. Psychopharmacology (Berl) 187:143–153.

Fields HL, Margolis EB (2015) Understanding opioid reward. Trends Neurosci 38:217–225.

Garritsen O, van Battum EY, Grossouw LM, Pasterkamp RJ (2023) Development, wiring and function of dopamine neuron subtypes. Nat Rev Neurosci 24:134–152.

Gerrits MA, Petromilli P, Westenberg HG, Di Chiara G, van Ree JM (2002) Decrease in basal dopamine levels in the nucleus accumbens shell during daily drug-seeking behaviour in rats. Brain Res 924:141–150.

He L, Gooding SW, Lewis E, Felth LC, Gaur A, Whistler JL (2021) Pharmacological and genetic manipulations at the micro-opioid receptor reveal arrestin-3 engagement limits analgesic tolerance and does not exacerbate respiratory depression in mice. Neuropsychopharmacology 46:2241–2249.

Ikemoto S (2007) Dopamine reward circuitry: two projection systems from the ventral midbrain to the nucleus accumbens-olfactory tubercle complex. Brain Res Rev 56:27–78.

Jalabert M, Bourdy R, Courtin J, Veinante P, Manzoni OJ, Barrot M, Georges F (2011) Neuronal circuits underlying acute morphine action on dopamine neurons. Proc Natl Acad Sci U S A 108:16446–16450.

Johnson SW, North RA (1992) Opioids excite dopamine neurons by hyperpolarization of local interneurons. J Neurosci 12:483–488.

Keiflin R, Janak PH (2015) Dopamine Prediction Errors in Reward Learning and Addiction: From Theory to Neural Circuitry. Neuron 88:247–263.

Koob GF, Sanna PP, Bloom FE (1998) Neuroscience of Addiction. Neuron 21:467–476.

Lammel S, Lim BK, Malenka RC (2014) Reward and aversion in a heterogeneous midbrain dopamine system. Neuropharmacology 76 Pt B:351–359.

Lammel S, Ion DI, Roeper J, Malenka RC (2011) Projection-specific modulation of dopamine neuron synapses by aversive and rewarding stimuli. Neuron 70:855–862.

Lammel S, Hetzel A, Hackel O, Jones I, Liss B, Roeper J (2008) Unique properties of mesoprefrontal neurons within a dual mesocorticolimbic dopamine system. Neuron 57:760–773.

Lammel S, Lim BK, Ran C, Huang KW, Betley MJ, Tye KM, Deisseroth K, Malenka RC (2012) Input-specific control of reward and aversion in the ventral tegmental area. Nature 491:212–217.

Lecca D, Valentini V, Cacciapaglia F, Acquas E, Di Chiara G (2007) Reciprocal effects of response contingent and noncontingent intravenous heroin on in vivo nucleus accumbens shell versus core dopamine in the rat: a repeated sampling microdialysis study. Psychopharmacology (Berl) 194:103–116.

Lefevre EM, Pisansky MT, Toddes C, Baruffaldi F, Pravetoni M, Tian L, Kono TJY, Rothwell PE (2020) Interruption of continuous opioid exposure exacerbates drug-evoked adaptations in the mesolimbic dopamine system. Neuropsychopharmacology 45:1781–1792.

Liu C, Goel P, Kaeser PS (2021) Spatial and temporal scales of dopamine transmission.Nat Rev Neurosci 22:345–358.

Luscher C (2016) The Emergence of a Circuit Model for Addiction. Annu Rev Neurosci 39:257–276.

Madhavan A, Bonci A, Whistler JL (2010a) Opioid-Induced GABA potentiation after chronic morphine attenuates the rewarding effects of opioids in the ventral tegmental area. J Neurosci 30:14029–14035.

Madhavan A, He L, Stuber GD, Bonci A, Whistler JL (2010b) mu-Opioid receptor endocytosis prevents adaptations in ventral tegmental area GABA transmission induced during naloxone-precipitated morphine withdrawal. J Neurosci 30:3276–3286.

Martianova E, Aronson S, Proulx CD (2019) Multi-Fiber Photometry to Record Neural Activity in Freely-Moving Animals. J Vis Exp.

Matsui A, Williams JT (2011) Opioid-sensitive GABA inputs from rostromedial tegmental nucleus synapse onto midbrain dopamine neurons. J Neurosci 31:17729–17735.

Matsui A, Jarvie BC, Robinson BG, Hentges ST, Williams JT (2014) Separate GABA afferents to dopamine neurons mediate acute action of opioids, development of tolerance, and expression of withdrawal. Neuron 82:1346–1356.

Mazei-Robison MS et al. (2011) Role for mTOR signaling and neuronal activity in morphine-induced adaptations in ventral tegmental area dopamine neurons. Neuron 72:977–990.

McClain SP, Ma X, Johnson DA, Johnson CA, Layden AE, Yung JC, Lubejko ST, Livrizzi G, He XJ, Zhou J, Chang-Weinberg J, Ventriglia E, Rizzo A, Levinstein M, Gomez JL, Bonaventura J, Michaelides M, Banghart MR (2023) In vivo photopharmacology with light-activated opioid drugs. Neuron 111:3926–3940 e3910.

Morales M, Margolis EB (2017) Ventral tegmental area: cellular heterogeneity, connectivity and behaviour. Nat Rev Neurosci 18:73–85.

Patriarchi T, Cho JR, Merten K, Howe MW, Marley A, Xiong WH, Folk RW, Broussard GJ, Liang R, Jang MJ, Zhong H, Dombeck D, von Zastrow M, Nimmerjahn A, Gradinaru V, Williams JT, Tian L (2018) Ultrafast neuronal imaging of dopamine dynamics with designed genetically encoded sensors. Science 360.

Pothos E, Rada P, Mark GP, Hoebel BG (1991) Dopamine microdialysis in the nucleus accumbens during acute and chronic morphine, naloxone-precipitated withdrawal and clonidine treatment. Brain Res 566:348–350.

Pupe S, Wallen-Mackenzie A (2015) Cre-driven optogenetics in the heterogeneous genetic panorama of the VTA. Trends Neurosci 38:375–386.

Simantov R (1993) Chronic morphine alters dopamine transporter density in the rat brain: possible role in the mechanism of drug addiction. Neurosci Lett 163:121–124.

Spanagel R, Almeida OF, Shippenberg TS (1993) Long lasting changes in morphine-induced mesolimbic dopamine release after chronic morphine exposure.Synapse 14:243–245.

Spencer MR, Miniño AM, Warner M (2022) Drug Overdose Deaths in the United States, 2001-2021. NCHS Data Brief, no 457.

Threlfell S, Lalic T, Platt NJ, Jennings KA, Deisseroth K, Cragg SJ (2012) Striatal dopamine release is triggered by synchronized activity in cholinergic interneurons. Neuron 75:58–64.

Yuan L, Dou YN, Sun YG (2019) Topography of Reward and Aversion Encoding in the Mesolimbic Dopaminergic System. J Neurosci 39:6472–6481.

